# Core Coefficient quantifies transcription-factor functional dominance for therapeutic target selection

**DOI:** 10.64898/2026.10.27.760230

**Authors:** Michael Klukinov, Rick Orr, William K. Martin, David C. Yeomans, Julien Mamet

**Affiliations:** Core Biotherapeutics, Inc., 2120 University Avenue, Berkeley, 94704, CA, USA

## Abstract

Transcription factors (TFs) operate in regulatory networks to control gene expression. The functional importance or leverage of a TF depends on the specific regulatory network in which it operates. Commonly used measurements of TF biology do not quantify the functional importance of TFs in these networks, which makes target selection for therapeutic intervention suboptimal. We developed the Core Coefficient (CC), a graph theory measure of TF functional dominance to overcome this limitation. Analysis of networks constructed from patient-biopsy and human post-mortem datasets revealed that these regulatory networks are organized in a functional hierarchy that is dominated by an identifiable and limited group of TFs, which we refer to as the Core. Analysis also revealed that the functional dominance of a TF, or the capacity to influence a cellular outcome, is a property of its position in the network hierarchy rather than its identity. While the Core consists of a group of functionally dominant TFs that control the network, the individual TFs forming that Core vary with cell type and cellular state. During the course of a given disease, from cancer to neurodegeneration, the functional dominance of TFs relevant to that disease shifts from low hierarchical importance to Core positions, whereas the functional dominance of other TFs is reduced or unchanged. CC can measure this shift in functional dominance and predict Core membership as a disease progresses. From a therapeutic standpoint, CC can also track the alterations in network hierarchy in diseased cells produced by treatments in the clinic. For example, network analyses of cancer cells before and after treatment from the same patients show dominant Core regulators shift from positions of high importance to positions of low importance after treatment. Experimental testing of these network mechanics in cancer cell lines showed CC can predict Core re-composition, signaling pathway reorganization and the cellular outcome of blocking a specific TF. In contrast, differential TF expression was informative in individual cases in these experiments but did not provide a general rule for measuring TF functional importance. This body of work introduces the CC, which quantifies TF functional dominance and enables the identification of targets for reprogramming TF networks to achieve a therapeutic objective.

## Introduction

TFs are a class of proteins that control fundamental cell properties and mechanisms, from cellular identity, differentiation and stress responses to proliferation and survival, through regulatory networks. These networks consist of individual TFs interacting with one another to collectively organize and regulate transcriptional programs.^1–4^ Depending on cell type and cellular state, an individual TF may occupy different positions within the overall regulatory TF network hierarchy. Consequently, a TF may perform distinct roles with varying degrees of functional importance, even when its DNA-binding and biochemical properties or expression level remain unchanged. Investigating TF biology therefore requires approaches that capture not only TF expression levels but also the functional importance of each TF within the regulatory network in which it operates.

Common approaches to studying TFs and nominating them as therapeutic targets evaluate them individually without accounting for the regulatory network in which they operate. Standard descriptive characteristics of TFs, such as differential expression, mutation, chromatin occupancy or genetic association, are informative and can link TFs to a phenotype, but none directly measure how much regulatory control a TF has over its network or what leverage it has over its cellular phenotype. This distinction becomes important when the objective changes from describing a biological state to deliberately altering its activity to achieve a therapeutic effect.

The development of the EGR1 TF decoy drug candidate known as AYX1 for postoperative pain illustrates this distinction. Despite strong evidence linking EGR1 to injury-induced neuronal plasticity and AYX1 demonstrating statistically significant treatment effects in multiple preclinical models of pain, together suggesting potential efficacy across a broad postoperative pain patient population, clinical benefit was observed only in a subgroup of patients with altered pain-regulation biology, known as catastrophizers.^5,6^ This example highlights the gap between establishing biological plausibility for a TF target and demonstrating that the TF occupies a functionally dominant, therapeutically actionable position within a disease-specific network and patient population.

Other drug candidates that targeted well-studied TFs but did not clearly deliver expected or repeatable clinical results over the past two decades include STAT3 inhibitors (approximately 130 clinical trials over 20 years with multiple therapeutic modalities produced mixed results and efficacy-safety imbalances), MYC inhibitors, and TEAD inhibitors^7–10^ The notable exception is belzutifan, sold under the brand name Welireg, an inhibitor of HIF2a that is approved for von Hippel–Lindau–associated renal cell carcinoma.^11^ Collectively, these cases underscore a clear discrepancy between the biological importance of TFs as a class of proteins and our current ability to identify the right TF or combination of TFs to target to achieve therapeutic benefits in a given disease or target patient population. As such, with over 1650 TFs encoded in the human genome^12^ and only a handful of TFs investigated clinically as targets, a novel approach to identifying TF targets with accurate prediction of therapeutic outcomes is needed to leverage the substantial therapeutic value of this class of proteins.

Human TF networks provide a basis for addressing this limitation in target identification. Previous studies showed that human TF networks share a common, modular architecture across cell types, despite being supported by distinct, cell-type-specific TF cohorts and regulatory interactions.^1–4^ This raises the possibility that, beyond network topology, TF systems also preserve a functional hierarchy in which the identities of TFs occupying particular hierarchical positions vary across cells, tissues, diseases, cellular states, and even patients.

We therefore asked two fundamental questions: first, can the functional dominance of individual TFs or groups of TFs within a network be measured quantitatively? Second, beneath the extensive cell- and state-specificity of TF regulatory networks, is there a conserved functional organization whose hierarchy or architecture persists even as the TFs occupying dominant positions change?

Answering these questions is important for understanding TF biology and therapeutic target identification for efficacy and tolerability. If regulatory control is functionally hierarchical, a TF-based therapeutic strategy should not rely only on which TF is expressed or differentially regulated. It should also ask which TF or group of TFs occupies a position with sufficient functional dominance to change the disease state when perturbed while being less dominant in normal cells to improve overall safety and tolerability of inhibiting the target TF.

To address these questions, we developed the Core Coefficient (CC), a graph-theory-based, systems-level measure of TF functional dominance within networks. We defined functional dominance as the capacity of a TF or group of TFs to influence, maintain, or reconfigure a network and its associated signaling or cellular state. With the CC, we first characterized the functional organization of networks constructed from healthy-subject samples and patient biopsy and post-mortem datasets spanning diverse cell types and tissues. These datasets represented mechanistically distinct diseases, including cancer, fibrosis, myocardial infarction, and neurodegeneration. We next examined networks across disease progression and paired pre- and post-treatment samples, and then directly tested the functional significance of the CC using independent cell-line experiments following TF blockade. These controlled perturbation experiments allowed us to determine how CC and functional dominance relate to cell signaling and cellular outcomes.

This novel approach to TF biology enabled us to characterize how TFs function within regulatory networks and validate CC as a framework for prioritizing TFs for therapeutic intervention and reprogramming dysregulated TF networks.

## Results

### CC delineates a hierarchical framework of TF functional dominance and identifies the network Core

The CC quantifies functional dominance and assigns each TF a corresponding value within its network. The absolute CC value is a quantitative measure of the functional dominance of each TF relative to other TFs in the same network. Within a given network, the distribution of absolute CC values indicates whether regulatory control is concentrated in a relatively small group of TFs or distributed more broadly across the network. Ranked CC converts absolute CC values into a numerical ordering, allowing TFs to be sequentially ranked according to their position within the functional hierarchy of a given network and facilitating efficient comparisons between networks. Together, these CC measures enable separating the common architecture of functional dominance from the identities of the TFs that instantiate it. In other words, CC measures show that functional dominance is a property of network position rather than identity.

To characterize the organization of TF functional dominance in human networks, we constructed networks from biopsy and post-mortem datasets comprising approximately 11,500 patients and 10 million single cells and analyzed them with the CC. The datasets included normal and diseased cells spanning more than 350 cell types and cellular states, as well as mechanistically distinct diseases, ranging from multiple cancer types and stages to fibrosis, myocardial infarction, and neurodegeneration.

Analysis of these networks showed that absolute CC values formed a non-uniform hierarchy. Most TFs occupied positions of low or intermediate dominance, whereas a small fraction exhibited disproportionately high dominance. The distribution of absolute CC values followed a two-segment structure, consisting of a steep upper-end segment of the most dominant TFs and a shallower segment spanning TFs with low to intermediate CC values. Piecewise regression identified the breakpoint at which the steep upper-end segment transitioned to the shallower distribution characterizing the remainder of the network (Figure 1a,b). In 87.5% of networks, the estimated breakpoint fell within the top 5% of TFs with the highest CC values. We therefore adopted the top 5% as a conservative, standardized operational definition of the regulatory Core, or the group of functionally dominant TFs that control a given network.

**Fig. 1.**
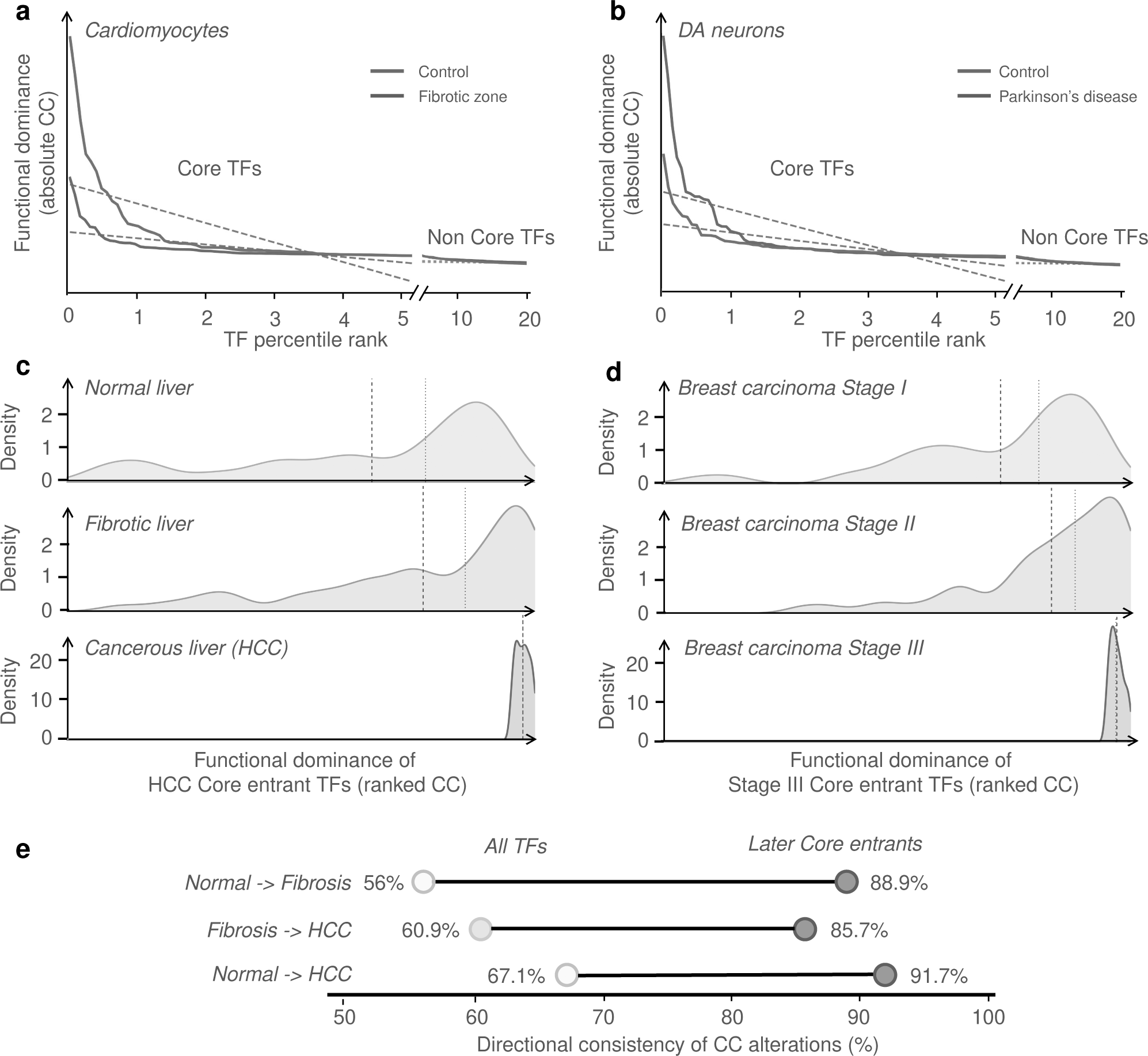
Disease-associated changes in TF network hierarchy architecture and Core composition. a,b, Examples of opposite changes in TF network hierarchy architecture between control and disease states, calculated using absolute CC. TFs were sorted by CC values, and the Core was defined as the top 5% of TFs. a, In heart cardiomyocytes, the Core distribution became steeper from control to the infarct fibrotic zone after a myocardial infarction, whereas the slope of the non-Core distribution changed little. b, The opposite behavior was observed in dopaminergic (DA) neurons in Parkinson disease, where the Core distribution became flatter relative to control DA neurons while the non-Core distribution remained comparatively stable. Solid lines show the observed absolute CC distributions; dashed lines show linear fits used to quantify Core and non-Core slopes of CC distribution. c, Progressive increase in ranked CC across normal liver, fibrosis and HCC. Kernel density estimates show the distributions of ranked CC values for the 56 TFs belonging to the HCC Core with measurable CC in the normal liver network. The same TFs were evaluated across all three states. Dashed and dotted vertical lines indicate the mean and median, respectively. Differences in CC values across disease states were significant (Friedman test, p = 4.18 × 10 ^−16^, n = 56). All three pairwise comparisons were significant (paired Wilcoxon signed-rank test with Holm correction): normal vs fibrosis (unadjusted p = 0.01376; adjusted p = 0.01376), fibrosis vs HCC (unadjusted p = 9.01 × 10^−1^; adjusted p = 1.80 × 10 ^−9^), and normal vs HCC (unadjusted p = 7.55 × 10^− 11^; adjusted p = 2.26 × 10^−1^). d, Progressive increase in ranked CC across breast cancer stages I, II and III. Kernel density estimates show the distributions of ranked CC values for 34 TFs belonging to the Stage III Core and measurable in the Stage I network. The same TFs were evaluated across all three stages. Dashed and dotted vertical lines indicate the mean and median, respectively. Overall differences were significant (Friedman test, p = 2.32 × 10^−9^, n = 34). All three pairwise comparisons remained significant after Holm correction (Stage I vs II, p = 7.42 × 10^−5^; Stage II vs III, p = 8.12 × 10^−9^; Stage I vs III, p = 3.49 × 10 ^−1^). e, Directional consistency of CC increases during liver disease progression. Paired-dot plots show the percentage of all TFs and later-state Core entrants whose CC changes were consistent with the direction of the indicated transition (Normal → Fibrosis, Fibrosis → HCC, or Normal → HCC). Percentages are displayed above each point. Later-state Core entrants show greater directional consistency than the overall TF population across all three transitions. Graph units are arbitrary unless specified.

Analysis of the relationship between CC values and the identity of the individual TFs that make up the network showed that the CC hierarchy was not fixed but instead contained both flexible and constrained positions. Across networks reconstructed from independent samples within the same broad biological state, individual TFs could occupy different positions in the CC hierarchy, indicating that similar cellular states can be supported by alternative TF-network configurations. This is particularly evident for TFs with mid-to-low CC values.

In contrast, TFs associated with reproducible biological transitions such as the shift from normal to diseased state or the progression across disease stages showed more consistent behavior. Their CC values were more likely to change concordantly across sequential biological states, increasing or decreasing in the same direction at each transition. This directional consistency was strongest among TFs that either (a) left the Core of the initial or early-stage network or (b) entered the Core at more advanced transition stages, such as later-stage disease or end-state cellular differentiation. Thus, whereas the broader TF network exhibits substantial configurational variability, TFs that become functionally dominant and acquire high CC values during biological state change follow more reproducible, directional patterns.

### Biological state change directionally recruits TFs into dominant Core positions in a programmed manner

To characterize changes in TF functional dominance across biological states and disease stages, we first analyzed liver diseases as they provide a unique longitudinal setting in which normal tissue progresses to fibrosis and subsequently to hepatocellular carcinoma (HCC). Because fibrosis and HCC represent mechanistically distinct diseases, this framework enabled us to track how TF dominance evolves across sequential pathological transitions and to identify regulatory features that reflect deeper principles of network reorganization beyond disease-specific phenotypes. The liver CC-network data show that later-state Core TFs do not appear at dominant positions randomly. TFs entering the HCC Core were already moving toward higher functional dominance in the preceding fibrotic state (Figure 1c). Across all TFs from liver networks, 56.0% showed increased CC from normal liver to fibrosis, 60.9% from fibrosis to HCC, and 67.1% from normal liver to HCC. In contrast, among TFs recruited to the Core of the later state, the corresponding proportions were 88.9%, 85.7%, and 91.7%, respectively. Thus, TFs recruited to the fibrosis or HCC Core were substantially more likely than the broader network background to show increasing CC during the transition to these states (Figure 1e). Notably, TFs occupying the fibrosis or HCC Core were not recruited from the network at random: they were already preferentially shifting toward greater functional dominance in the preceding disease transition. These findings support a model in which Core recruitment during disease progression reflects a directional reorganization of the TF regulatory network rather than a stochastic redistribution of TF dominance.

The same principle of progressive acquisition of later-stage Core positions appeared in other settings. Analysis of network hierarchy evolution through progression of independent cancer types showed that a similar pattern was observed across the progression of invasive breast carcinoma from Stage I to Stage II to Stage III (Figure 1d). TFs that were specific to the Stage III Core progressively increased in ranked CC across the earlier stages, with mean ranked CC increasing by 20% from Stage I to Stage II and by ∼10% in Stage III. In contrast, TFs specific to the Stage I Core progressively lost dominance across the same sequence. Similarly, in networks from bladder urothelial carcinoma, TFs specific to the Stage IV Core progressively increased in mean ranked CC by ∼10% from Stage II to Stage III and by ∼25% to Stage IV, while TFs specific to the Stage II Core progressively lost dominance. A similarly ordered trajectory was observed in stomach adenocarcinoma, where Stage III Core-specific TFs increased by ∼45% from Stage I to Stage II and by ∼20% to Stage III, while Stage I Core-specific TFs decreased by ∼10 and ∼25% through the same stages, respectively. On average, the CC increase from early (I-II) to late (III-IV) cancer stages was 22 ± 4%, p = 0.03 (Student’s t-test).

The other tumor types we analyzed included colon adenocarcinoma, lung squamous cell carcinoma, thyroid carcinoma, and kidney renal clear cell carcinoma. Networks of these tumor cells showed the same organized gain and loss of functional dominance as disease progressed through stages but followed different intermediate trajectories. This indicates that disease progression is programmed from a TF dominance standpoint and follows a state-specific program of TF recruitment into, and loss from, Core positions.

In summary, the movement of TFs that comprise the Core of late- and end-stage disease is predictable and detectable across earlier disease stages, before those TFs ultimately enter a position of dominance in the Core of the end-stage disease.

Beyond cancer, myocardial infarction networks showed a similar progressive reorganization of CC toward the most affected disease state. In the dataset we analyzed, disease-associated changes could be examined spatially rather than temporally, with the hierarchy of cardiomyocyte and fibroblast networks progressively changing from control tissue through the remote zone to the actively remodeling border zone of the infarct. Post-mortem neurodegenerative datasets from Alzheimer’s, Parkinson’s and Huntington’s diseases and iPSC-to-motor-neuron differentiation, as an experimental model of developmental network mechanics, provided additional demonstrations in which Core TF occupancy changed progressively in a state-specific manner while the higher-order organization of CC was retained.

### CC predicts Core membership recruitment and tracks therapeutic treatment effects

CC contains predictive information about later Core membership, starting at early stages of disease progression. In liver diseases, fibrosis-state ranked CC predicted subsequent HCC Core membership with a receiver-operating-characteristic area under the curve (ROC AUC) of approximately 0.83, where 0.5 indicates random discrimination and 1.0 perfect discrimination. Similarly, in breast carcinoma, Stage II CC predicted Stage III Core membership (ROC AUC ≈ 0.92). Adding the change in ranked CC from Stage I to Stage II did not improve prediction beyond Stage II CC alone. This suggests that a TF’s current position in the regulatory hierarchy captures more information about its subsequent Core membership than its preceding change in dominance. Notably, predictive information was already present at Stage I across the ten cancer types we analyzed: HCC, breast invasive carcinoma, colon adenocarcinoma, lung adenocarcinoma, lung squamous cell carcinoma, melanoma, uterine corpus endometrial carcinoma, thyroid carcinoma, kidney renal clear cell carcinoma, and stomach adenocarcinoma. Across these cancers, Stage I ranked CC predicted Stage III Core membership with ROC AUC values ranging from 0.70 to 0.89. Together, these results show that CC can identify TFs positioned to acquire regulatory dominance well before they enter the later-state Core, providing a unique capability for therapeutic target selection during disease progression.

From a therapeutic standpoint, successful treatments can profoundly alter the population of TFs that make up the Core. In paired, breast-tumor samples pre- and post-treatment from patients treated with independent therapy (in this case, clinical trials with patients with ER^+^ or HER2^+^ tumors treated with hormonal or chemotherapy combined with a targeted therapy), Core composition was markedly reorganized after treatment. This reorganization included loss of established oncogenic TFs, including FOXM1 and DNMT1 in ER^+^ tumors and HMGA2 and USF2 in HER2^+^ tumors, together with recruitment of new dominant TFs with established tumor-suppressive activity in breast cancer, including FOXN2 and ZBTB4 in ER^+^ tumors and RFX1 in HER2^+^ tumors.^13–19^ Together, these findings show that CC not only anticipates future Core membership but also captures the dynamic functional reorganization of TF dominance that accompanies therapeutic response.

### Experimental TF blockade validates CC as a measure of functional dominance

We next tested experimentally whether CC predicted the consequence of TF blockade on cellular outcome. The network of hard-to-treat MDA-MB-231 triple-negative breast cancer cells was built from a large number of independent datasets and analyzed with the CC. Based on this analysis, oncogenic TFs associated with MDA-MB-231 and closely related cells were selected to represent targets with a range of CC values, from low to high, and corresponding TF decoy inhibitors were designed (see Materials and Methods). Cells were transfected with vehicle or the decoy inhibitors and monitored every 3 hours for confluence and apoptosis using a Caspase-3/7 dye with an Incucyte Live-Cell Analysis System over 120 hours. The confluence of untreated control cells increased ∼ 4-fold over the 120h experiment (Figure 2a), showing robust cellular growth. Cell transfection with a reference decoy conjugated to Alexa Fluor 488 confirmed decoy transfection and showed it peaked ∼ 15h after transfection initiation (Figure 2b). Growth responses differed significantly among the RNAiMAX control and the D231-1, D231-6, and D231-8 decoy treatments (Welch’s one-way ANOVA, p = 0.0026, n = 9) (Figure 2a). Analysis of the entire cohort of tested decoys shows that higher CC values of decoy TF targets were strongly associated with lower cell confluence after blockade (Spearman ρ =-0.949, p = 9.2 × 10^−5^, n=9) and greater confluence-normalized Caspase-3/7 burden (ρ =0.797, p=0.0102, n = 9) (Figure 2c,d). Apoptotic burden was itself associated with reduced cell expansion (ρ =-0.86, p=0.014, n = 9). Thus, in this experiment, functional dominance measured by the CC quantitatively and precisely tracked the subsequent biological consequence of blocking targeted oncogenic TFs. Together, these results experimentally validate CC as a quantitative predictor of TF dependency, with higher CC values identifying TFs whose blockade more effectively suppresses cancer cell expansion and induces their apoptosis.

**Fig. 2.**
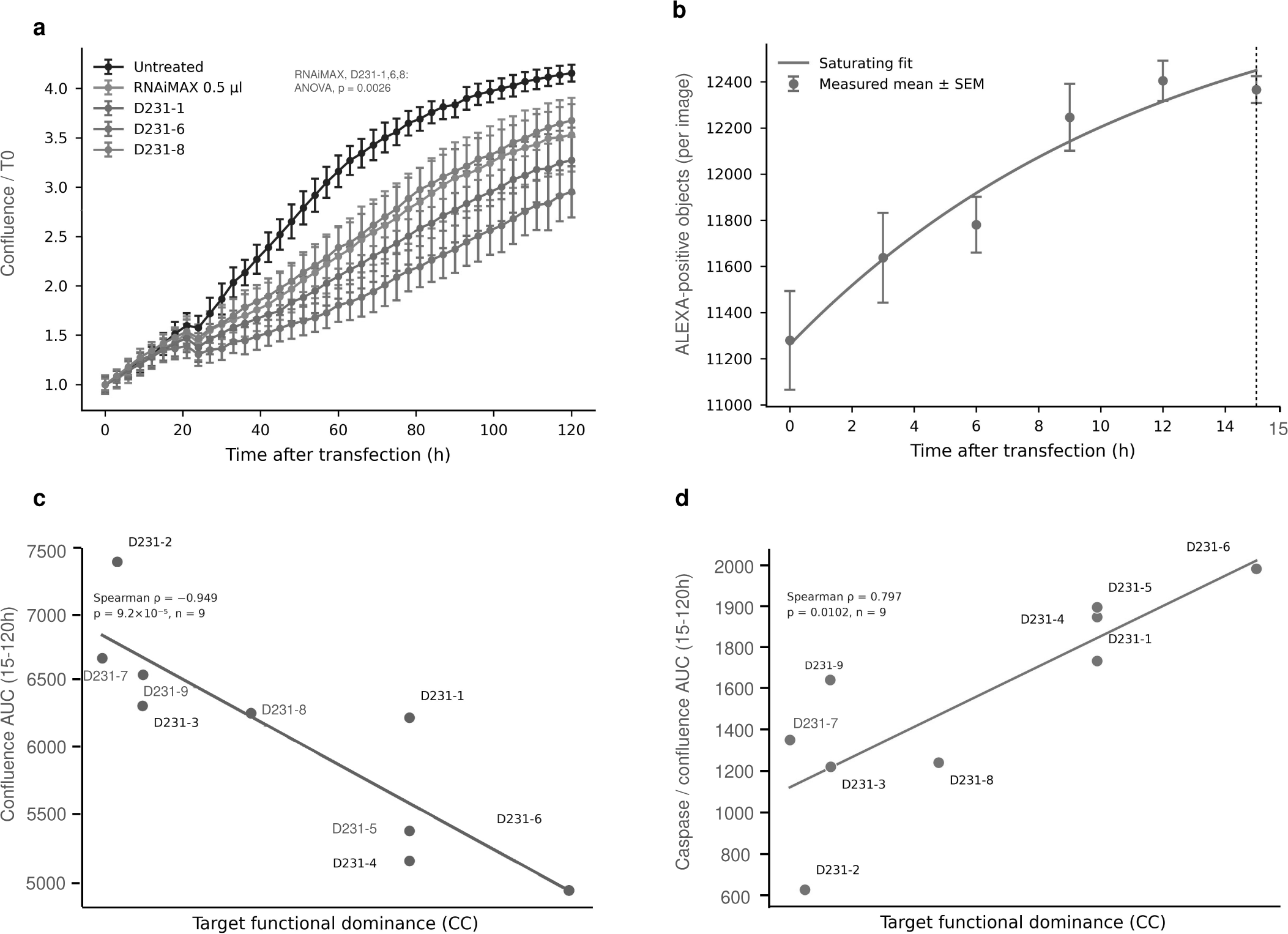
CC predicts the cellular response to oncogenic TF blockade in MDA-MB-231 cells. a, Confluence of MDA-MB-231 untreated cells (vehicle control) and treated with RNAiMAX (transfectant control) or TF decoys. Confluence was normalized to the value measured at the time of plating and the start of imaging (T0). Data are presented as mean ± SEM of each measure made during the 120h-long experiment. Points show measured means and error bars indicate SEM. The area under the cell-confluence curve (AUC) from 15h to 120h, spanning maximal transfection efficiency to the end of the experiment, differed significantly among the RNAiMAX transfectant control, D231-1, D231-6, and D231-8. RNAiMAX is used as a control comparator as it produces a baseline toxicity over cells as compared with the vehicle treatment. b, Transfection kinetics measured using Alexa488-conjugated decoy over the first 15 h after transfection. Points show mean Alexa488-positive objects per image ± SEM. The curve shows the saturating fit to the measured data; the dashed vertical line indicates 15 h, the beginning of the interval used for subsequent AUC analyses as it marks the time of maximum transfection. c, Relationship between target TF functional dominance, measured by CC, and confluence AUC from 15h-120h for the nine tested decoys. Greater functional dominance of the targeted TFs was associated with lower cell confluence following decoy blockade. d, Relationship between target TF functional dominance and caspase signal normalized to confluence over the same 15h to 120h interval. Greater functional dominance of the targeted TFs was associated with greater apoptotic activity following decoy blockade. Graph units are arbitrary unless specified.

### Differential expression is informative in selected cases but is not a general rule for TF functional importance

We next tested a common assumption used in TF target nomination or assignment of importance in a given cell type, tissue or disease: if expression of a TF increases as a cellular state develops, that TF is becoming more functionally important, whereas decreasing expression implies reduced importance. We tested this directly in MCF7 ER^+^ cells, another breast cancer model that is well described and practical to use. We selected 12 TF targets with known oncogenic roles and designed decoy inhibitors against them (see Materials and Methods). Cells were plated, transfected with vehicle or decoys and grown for either 4h, a baseline timepoint that enables cells to settle after plating and transfection, or 72h, a time point that enabled a 2.5-fold increase in cell number from baseline in our experimental conditions. Transfection was confirmed at 24h by FACS using a fluorescein-conjugated decoy. Measures of total cell count and number of cells per cell cycle phase (G1, S, G2/M) as well as CellTiter-Glo reading of cell viability were performed at both timepoints. Gene expression was also quantified, and TF networks were constructed and analyzed using CC. Reactome signaling pathway scores were also calculated for each testing condition.

Differential expression of the targeted TFs between 4 and 72h in vehicle-treated cells did not predict the effects of the corresponding decoy treatments on cancer cell numbers (Spearman ρ = 0.224, p = 0.484, n = 12) or CellTiter-Glo signal (ρ = -0.119, p = 0.713, n = 12) (Figure 3a,b). Differential TF expression did not predict cell-cycle redistribution, Core re-composition or global Reactome-program reorganization either. Separate comparisons within each cell cycle phase gave the same overall result. Expression can nevertheless agree with biological consequence for some individual TFs and can associate with selected network properties but was limited to a case-by-case basis (Figure 3a,c, see XBP1 TF). This does not imply that expression is uninformative. Rather, differential expression provides biological context, but these data support the conclusion that differential expression of a TF on its own is not sumcient as a general measure of TF functional importance.

**Fig. 3.**
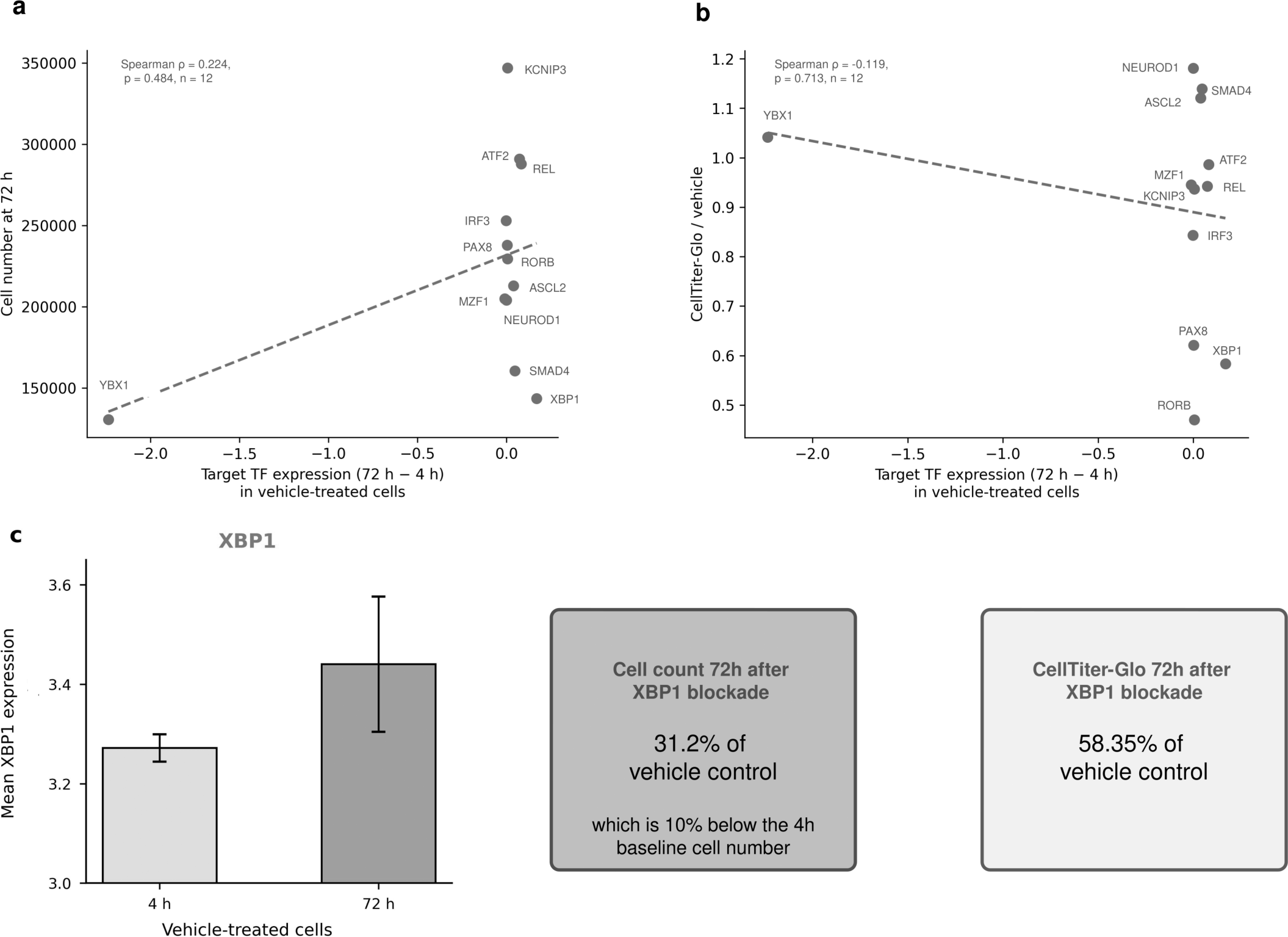
Changes in target TF expression do not consistently predict the cellular response to TF blockade in MCF7 cells. For each TF, expression change was calculated as its mean expression at 72h minus its mean expression at 4h in vehicle-treated cells. Positive values indicate an increase in expression from 4h to 72h, whereas negative values indicate a decrease. a, Relationship between the change in target TF expression level in vehicle-treated cells from 4h to 72h and cell number at 72h following blockade of the corresponding TF. The isolated point at the left is separated by an axis break to display the remaining targets at an informative scale. b, Relationship between the same change in target TF expression and CellTiter-Glo signal at 72h following target blockade, expressed relative to the corresponding vehicle-treated control. c, XBP1 as an example of a TF whose expression increased between 4h and 72h in vehicle-treated cells. Bars show mean XBP1 expression across G1, S and G2/M ± SEM at each time point. The corresponding cellular outcomes following XBP1 blockade are shown at right: cell number fell below the experimental baseline, and CellTiter-Glo was reduced relative to the vehicle control. Together, these analyses illustrate that changes in TF abundance can accompany sensitivity to blockade in individual cases but do not provide a general predictor of cellular response across targets. Graph units are arbitrary unless specified.

### MCF7 perturbations show how functional dominance propagates through the network

We used the same MCF7 ER^+^ cell experiment to resolve how network dominance is altered after selective TF blockade. G1-, S-and G2/M-phase-specific networks were constructed from vehicle-and decoy-treated cells. There are approximately 1,650 human TFs, which can be grouped into about 100 families, allowing us to examine the role of functional dominance in network reorganization at both the individual TF and TF family levels.

Analysis of untreated (i.e., vehicle control) MCF7 networks showed that individual TF dominance changed substantially across cell-cycle phases, whereas the dominance of TF families, computed as the sum of the CC values of the TF family members, remained comparatively stable. Spearman rank correlations of family CC between cell-cycle phases ranged from approximately 0.8 to 0.9, indicating that family-level regulatory organization can persist while individual TFs exchange dominance.

We next examined whether baseline functional dominance of individual TFs predicted changes in Core composition following TF blockade. We compared Core membership between samples collected 4 and 72 hours after treatment with vehicle or TF decoys. To ensure comparability, Core disturbance analyses were restricted to the 10 TFs represented in both the 4h and 72h G1 networks, with G1 used as the baseline cell-cycle state. As such, two TFs did not meet the predefined inclusion criteria for this analysis: ASCL2 because post-treatment network data at 72h could not be generated and NEUROD1 because it did not meet the required G1 network-representation criterion at 4h and 72h. Target CC was defined as the CC value of the target TF. Higher Target CC in untreated G1 networks at 4h was strongly associated with greater G1 Core disturbance at 72h (Spearman ρ = 0.685, p = 0.0289, n = 10; Figure 4a). The observed correlation remained positive but was moderate for the subsequent S phase (ρ = 0.491, p = 0.149; n = 10; Figure 4b) and G2/M (ρ = 0.467, p = 0.173, n = 10; Figure 4c). These observations suggest that the consequences of targeting a TF with high functional dominance in G1 can extend across cell-cycle phases, although the strongest association was observed within G1. In contrast, baseline G1 target expression for the same 10 TFs showed a modest, negative non-significant association with subsequent G1 Core disturbance (ρ = –0.37, p = 0.293, n = 10) (Figure 4f), indicating no evidence of the positive predictive relationship observed with the CC. Thus, baseline TF functional dominance provided information about subsequent Core reorganization that TF expression did not capture.

**Fig. 4.**
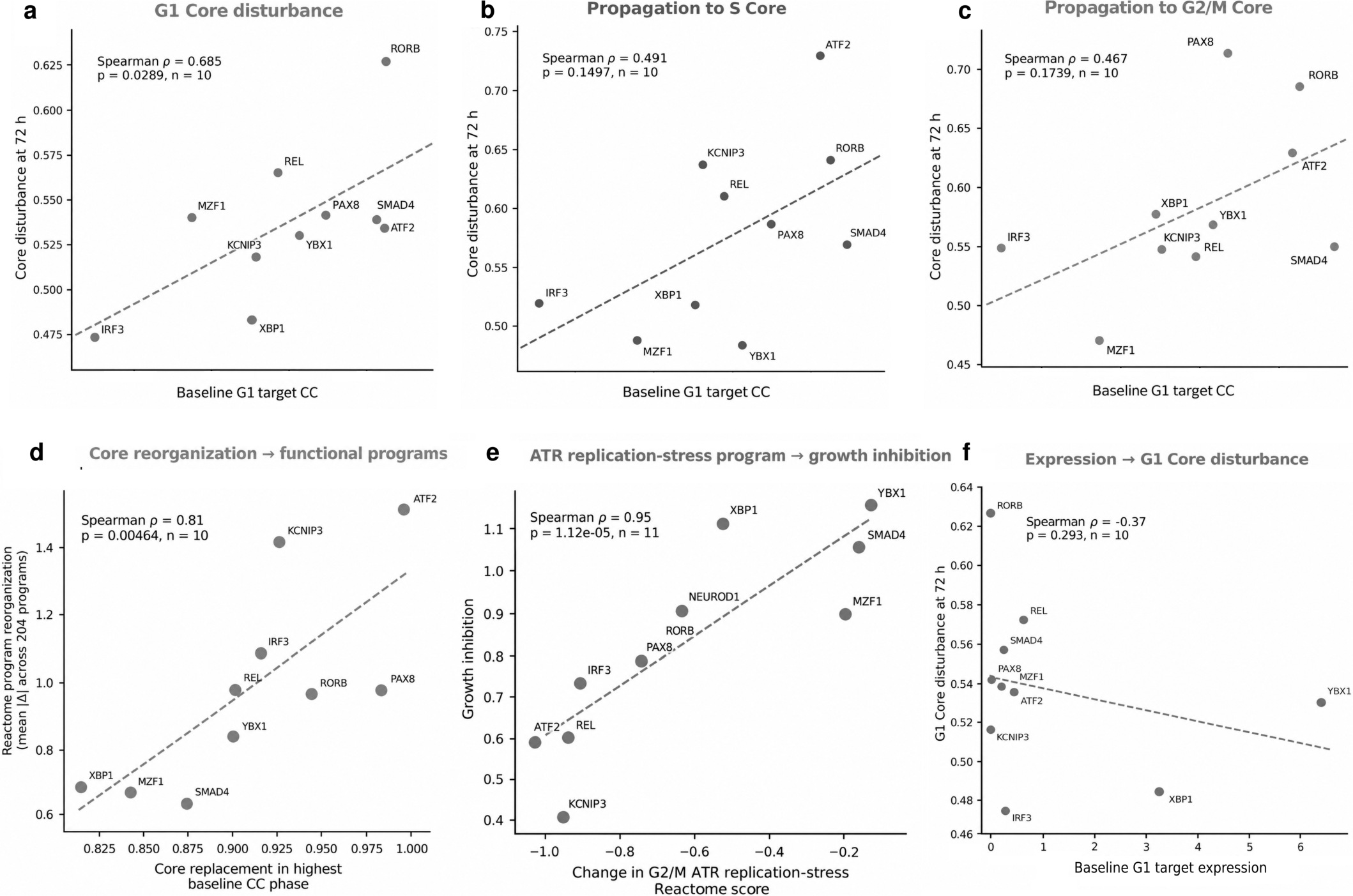
CC identifies state-specific Core disruption linked to functional-program reorganization and cellular outcome. a–c, Relationship between TF target functional dominance at baseline and subsequent Core reorganization following TF blockade. Target CC was measured in G1 in vehicle-treated cells at 4h. Core disturbance was calculated as the average of the fractions of TFs lost from the untreated Core and newly recruited into the treated Core, counting only changes exceeding untreated variability. a, Higher baseline G1 Target CC was associated with greater G1 Core disturbance at 72h. b, This relationship propagated into S phase, although it was weaker. c, A similar relationship was observed in G2/M. d, Relationship between Core replacement and functional-program reorganization. For each perturbation, the 72h cell-cycle phase was selected according to the phase in which the targeted TF had its highest CC in vehicle-treated cells at the 4h baseline. Reactome reorganization was calculated as the mean absolute change across 204 Reactome program scores between decoy- and vehicle-treated cells in the same phase. Greater Core replacement was associated with greater Reactome program reorganization. e, Relationship between reorganization of the ATR replication-stress program and cellular outcome. The magnitude of change in the G2/M Reactome program Activation of ATR in response to replication stress was strongly associated with growth inhibition across TF perturbations. Growth inhibition was calculated as 1 minus the ratio of net cell growth from 4h to 72h in decoy-treated cells to that in vehicle-treated controls. A value of 0 indicates no growth inhibition, whereas a value of 1 indicates complete suppression of cell growth. f, Baseline G1 target expression was tested against the same 10 G1 Core-disturbance values from panel a. Baseline G1 target expression was not associated with subsequent G1 Core disturbance. Graph units are arbitrary unless specified.

For some TF targets, including IRF3 and ATF2, the largest change in Core composition following their blockade occurred during the cell-cycle phase in which the targeted TF had the highest baseline CC. For others, including REL and RORB, the greatest Core change emerged during a later phase. Thus, CC identified the cell-cycle state in which a TF had the greatest functional leverage, whereas the resulting reorganization of the network could become apparent either in that same state or later.

We then asked whether the organization of functional dominance within TF families affected the network response to blockade of a single TF. Target Family Leverage was defined as Target CC divided by the CC of its TF family before treatment. For comparison with the analyses of individual TF CC, Target Family Leverage was calculated for the same 10 TFs as above. We examined Target Family Leverage separately in G1, S and G2/M. The highest observed correlation between Target Family Leverage and subsequent whole-network reorganization was in G1 (Spearman ρ = 0.733, p = 0.0202, n = 10), whereas the correlations were weaker in S (ρ = 0.261, p = 0.4697, n =10) and G2/M (ρ = 0.370, p = 0.2957, n =10). Target Family Leverage was redistributed substantially within several TF families across cell-cycle phases. For example, IRF3 carried 3% of its family CC in G1 but 45% in G2/M, whereas ATF2 fell from 16% to less than 1%. Thus, the functional importance of a TF cannot be considered independently of either its TF family or the cellular state in which that family operates.

To assess whether the G1 association depended on any individual TF, we performed a leave-one-target-out sensitivity analysis across all ten targets. The association between baseline G1 Target Family Leverage at 4h and subsequent whole-network reorganization following TF blockade at 72h remained positive in every iteration (Spearman ρ = 0.633–0.883, p = 0.0031–0.0760), with six of ten iterations reaching nominal statistical significance. These findings indicate that the positive relationship between Target Family Leverage and network reorganization was not driven by any single TF, supporting the role of family-relative functional dominance in shaping network responses to TF blockade. In biological terms, blocking a TF is associated with greater network reorganization when that TF carries a larger share of its family’s functional dominance, consistent with reduced potential for compensation by related TFs.

### Core replacement is accompanied by functional-program and cellular change

We next asked whether Core reorganization was accompanied by changes in biological programs. For each MCF7 decoy perturbation described above, we compared scores for 204 Reactome pathways^20^ between vehicle- and decoy-treated cells in the same cell-cycle phase. We used the magnitude of the score change to measure pathway reorganization. Because the direction of these scores does not by itself establish pathway activation or repression, absolute score changes were analyzed.

A clear correlation between CC and biological programs emerged when each perturbation at 72h was examined in the cell-cycle phase in which the target had its highest CC in vehicle-treated cells at 4h. Two TFs did not meet predefined inclusion criteria for this analysis: ASCL2, because 72h treated data could not be generated as mentioned above, and NEUROD1 because its CC was null across all cell-cycle phases at the 4h baseline, preventing selection of a highest-CC phase. Across 10 TF perturbations, greater replacement of the untreated regulatory Core in the phase of highest baseline Target CC was associated with greater Reactome program reorganization (Spearman ρ=0.81, p=0.0046, n = 10) (Figure 4d). In contrast to CC, selecting the phase in which the same target TF had its highest baseline expression did not recover this relationship (Spearman ρ=0.03, p=0.934, n =10). Thus, CC identified the cellular state in which disruption of the network Core was coupled to broader functional-program reorganization, whereas target expression did not provide the same state information. Hence, in this systematic TF blockade experiment, CC provided information about functional control over signaling pathways that transcript abundance alone did not provide.

The altered programs also matched cellular outcome. In G2/M, change in a kinetochore/chromosome-segregation program tracked the fraction of cells in G2/M. The Ataxia-Telangiectasia and Rad3-related (ATR) replication-stress program was strongly associated with growth inhibition (Spearman ρ = 0.95, p = 1.12 × 10^−5^, n = 11) (Figure 4e). These examples connect TF blockade to Core replacement, functional-program change and, where an appropriate phenotype was measured, altered cell behavior.

Changes in Core composition were also associated with changes in the proportion of cells in each cell-cycle phase following TF decoy blockade. Greater replacement of the G1 Core was associated with a lower proportion of cells in G1 and a higher proportion in S phase. The cellular effect, however, depended on which TF was blocked. For example, XBP1 blockade reduced cell numbers below the 4h baseline and reduced the CellTiter-Glo viability signal relative to the vehicle-treated control, consistent with loss of cell growth or survival. Other targets produced changes in the proportion of cells in G1, S or G2/M without the same loss of the cell population when blocked. Thus, Core replacement measures how strongly the dominant regulatory program is reorganized, but it does not specify the cellular response. For this, CC needs to be paired with the known function of the TF being blocked to determine whether that reorganization is expressed as altered cell-cycle progression, reduced growth, or cell loss.

Together, experimental data support a layered mechanism of network reprogramming: (1) CC identifies which TF or group of TFs carries functional leverage in a given regulatory state; (2) blockade of those TFs changes the composition of the Core; (3) Core replacement is linked to functional-program reorganization; and (4) program change can be linked to a function-matched cellular outcome. Thus, these data support that the CC can predict sequential mechanistic associations relevant to the selection of targets to achieve desired therapeutic outcomes by reprogramming TF networks.

## Discussion

The central result of this body of work is that functional dominance is a measurable dimension of TF operations that presents its own biology. Patient- and tissue-derived networks repeatedly showed a hierarchy dominated by a Core of restricted size. The TFs occupying the different compartments of that hierarchical organization are context dependent, but their behavior is not arbitrary: TFs recruited to positions of dominance or dismissed to positions of low dominance during cellular or tissue state change were more directionally constrained than TFs sitting in the background of network dominance. In particular, TFs recruited into the Core at later stages of cellular or disease progression showed strong directional conservation across state transitions. The conserved property across cell types, tissues or diseases is therefore not a fixed TF list, but an organization of regulatory control in which different TFs can occupy equivalent functional positions depending on context.

This framework distinguishes the relationship between TF expression and functional dominance. Differential expression can identify important TFs and can agree with perturbational consequences in individual cases. Our experimental data show that this interpretation does not extend as a general rule across TFs: the direction of TF expression change did not consistently predict the regulatory or cellular response to TF blockade. CC addresses a different question than differential levels of expression by asking where functional leverage resides within the network. This is consistent with independent observations from the literature: for instance, induction of the FOXO3a tumor suppressor in MDA-MB-231 cells has little effect on cell proliferation and, similarly, although STAT5 is known to promote proliferation, its overexpression does not necessarily alter proliferative output.^21,22^ On the other hand, a low expression level can also mean increased dependence on a TF. That is the case for SPI1 in acute myeloid leukemia where SPI1 levels are down from baseline, but leukemic cells require its residual, low-level expression for survival.^23^ Expression and CC should therefore be viewed as complementary measurements rather than competing descriptions of the same variable.

Network perturbation experiments give functional significance to this framework. MDA-MB-231 experiments confirmed that higher predicted TF dominance was associated with stronger blockade consequences when TFs are working within the same biological function, in this case, oncogenicity. MCF7 experiments resolved the intervening sequence from functional dominance to Core reorganization, to broader program reorganization, and finally to a cellular phenotype consistent with the affected program. Family organization adds another dimension because the consequence of blocking one TF depends partly on how much of its family’s functional dominance it carries.

These findings provide a biological basis for changing the paradigm for TF target identification on a disease- and target patient population-, or even individual patient-specific basis. A key question when selecting a TF target is not only which TF changes in disease, but which TF has acquired enough functional leverage that inhibiting it can reorganize the disease state. A practical workflow is to reconstruct the relevant TF network, measure TF and family dominance, identify the Core, compare it with appropriate healthy or treated states, and prioritize TFs expected to alter the disease-associated regulatory program for therapeutic target selection.

## Conclusions

The results support a reproducible framework in which TF regulatory systems preserve a hierarchy of functional control over cells while exchanging TFs that occupy dominant positions across cells and cellular states. CC provides a quantitative way to identify those positions and to distinguish flexible background regulation from the directionally constrained regulators recruited during state change. This provides a framework for understanding TF function in network context and, from that biological foundation, a strategy for selecting TF targets intended to reprogram a pathological state toward cell death in the case of cancer cells, or restoration of normal regulatory programs in other cases such as neurodegeneration or post-infarction cardiomyopathy.

## Acknowledgments

The authors are grateful to the National Science Foundation for funding the MCF7 cell line work under Award No. 2451628. Any opinions, findings, conclusions, or recommendations expressed in this material are those of the authors and do not necessarily reflect the views of the National Science Foundation.

## Materials and Methods

### Human tissue and patient-sample sources for constructing TF networks

Networks were constructed from healthy subjects and patient biopsies or post-mortem datasets. Networks were analyzed within state, between individuals and, where biologically justified, across disease, spatial or differentiation states. Datasets used for the analyses presented in the publication:

- Cancer: TCGA (https://www.cancer.gov/ccg/research/genome-sequencing/tcga), METABRIC (https://ega-archive.org/studies/EGAS00000000083), breast cancer pre and post treatment (GSE153470, GSE119262, GSE87455)
- Alzheimer’s disease: GSE138852
- Parkinson disease: GSE178265, GSE157783
- Huntington’s disease: GSE138525, GSE180928, GSE233387, GSE270728, GSE152058, nemo:dat-ztfn3cc
- Liver, HCC: GSE33006, GSE25097, GSE104580, GSE101685, GSE29721, GSE102079, GSE69715, GSE116174, GSE107170, GSE112790, GSE99807, GSE22058, GSE17548; Liver, fibrosis: GSE19665, GSE45436, GSE121248, GSE9843, GSE62232; Liver, normal: GSE6764, GSE84402, GSE19665, GSE29721, GSE33006, GSE40873, GSE22058, GSE69715, GSE84402, GSE102079, GSE107170, GSE25097, GSE121248, GSE6764, GSE14668, GSE14951, GSE23343, GSE7117, GSE28619, GSE38941, GSE63067, GSE83148, GSE96851, GSE101685, GSE102079, GSE107037, GSE112790, GSE133815, GSE152738, GSE37031
- Human iPSCs and motor neurons: GSE138121
- MDA-MB-231: GSE152529, GSE156802, GSE52828, GSE137842, GSE110202, GSE98237, GSE113571, GSE71363, GSE49962, GSE83818, GSE67966, GSE45827, GSE59228, GSE59230, GSE59229, GSE56084, GSE36441, GSE63138, GSE50811, GSE60123, GSE60124, GSE60878, GSE60879, GSE57083, GSE35525, GSE51739, GSE46925, GSE23362, GSE40057, GSE44239, GSE41445, GSE39718, GSE39719, GSE39720, GSE32668, GSE33950, GSE39042, GSE33145, GSE36953, GSE33827, GSE35511, GSE32052, GSE22250, GSE24249, GSE30775, GSE25162, GSE29137, GSE26462, GSE21832, GSE19737, GSE15893, GSE18903, GSE18146, GSE11407, GSE13806, GSE15477, GSE15026, GSE14491, GSE10890, GSE12199, GSE11951, GSE10021, GSE9747, GSE9015, GSE20085, GSE36133, GSE66082, GSE137178, GSE121945
- Myocardial infarction: EGAS00001006330

### Network construction and Core Coefficient

The network construction and CC algorithms are proprietary, therefore numerical CC values are not provided on the figure axes.

### TF decoys

Decoys were manufactured by Integrated DNA Technologies, IA and resuspended in 10 mM Tris, pH 7.4 or DNase- and RNase-free water to prepare stock solutions. Sequences are provided for each TF in the next section below.

### MDA-MB-231 perturbation

MDA-MB-231 cells were transfected with vehicle or decoys using Lipofectamine RNAiMAX (Thermo Fisher Scientific, MA) in Opti-MEM and grown in DMEM (L-Gln, 1 g/l Glc, 1 mM NaPyruvate) with 10% HI FBS. TF decoys (target, 5’-3’ sense strand sequence) included single- and multi-target designs: D231-1, D231-4 and D231-5 were different sequences targeting the same TFs for assessing the effect of sequence on output (E2F factors, FOXC1, FOXM1, GTF2IRD1, TCF4, TGIF2 and ZEB2, AACAGAAGGTAAACAAAGATTATGGCGC-CTTTTGACAGGTG, AAAAGGCGCCATAATCTTTGTTTAC-CTTCTGTTGACAGGTG, CACCTGTCAACAGAAGGTAAA-CAAAGATTATGGCGCCAAA), D231-2 (FOXM1, ACA-GAAGGTAAACAAA), D231-3 (FOXC1, FOXM1 and GTF2IRD1, ACAGAAGGTAAACAAAGATTATGGC), D231-6 (CXXC5, FOXC1, FOXM1, MAX, NFIL3, TCF4, XBP1, YBX1, YY1 and ZEB2, CACCTGTTTATTTACACGTGT-TACATCACCGATGGAGCCATCTT), D231-7 (E2F factors, LYL1 and TCF7L1, GATCAAAGGAACAGATGGTTTTG-GCGCCAAAAG), D231-8 (GTF2IRD1, TCF4, TGIF2 and ZEB2, GTTGACAGCTGTCAGGATTAACGCACCTGCA), and D231-9 (FOXC1 and FOXM1, GAAACAAACAAACAATCGTAAATAAACAG). Decoy transfection data showed that transfection efficiency was highest at decoy concentration of 100 nM in that cell line. Confluence and Caspase-3/7-positive cells (detected as red objects with a red Caspase-3/7 dye (Sartorius, MI)) were measured every 3h with an Incucyte Live-Cell Analysis System (Sartorius, MI). Caspase counts were normalized to confluence. MDA-MB-231 CC values were calculated from an independent public transcriptomic dataset. Because target number correlates with CC in this small combinatorial series, the analysis is interpreted as functional validation of regulatory leverage rather than a clean single-target causal estimate. N = 3 per condition.

### MCF7 perturbation and expression analyses

MCF7 cells were transfected with vehicle or decoys using Lipofectamine RNAiMAX (Thermo Fisher Scientific, MA) in Opti-MEM and grown in EMEM with 0.01 mg/ml insulin and 20% FBS. Cells were reverse transfected with vehicle or decoys using Lipofectamine RNAiMAX (Thermo Fisher Scientific, MA) and grown over 72h. Decoys (target, 5’-3’ sense-strand sequence) were MCF7-H1 (MZF1, TGGGGATTGAGGGGGAA), MCF7-H2 (IRF3, GGAAAGGGAAACCGAAACCTGATTGGCT), MCF7-H3 (ATF2, CTGACATAAGATTACGTCAT), MCF7-H4 (REL, GGGAAATTCCCGGGGATTTCCC), MCF7-H5 (RORB, TAGGTCATGACCTACTGACCTA), MCF7-H6 (KCNIP3, GTCAACTGAGTCAACTGA), MCF7-H7 (PAX8, GGCTGAGCAGTCAAGCGTGACG), MCF7-H8 (NEUROD1, AACATATGTTGACATATGGC), MCF7-H9 (ASCL2, GCAGCTGCCGCGTCGTGT), MCF7-H10 (YBX1, GTACCATCGGCCAATCC), MCF7-H11 (SMAD4, CAGACAAGGTGCCAGACA) and MCF7-H12 (XBP1, TGACGTGGCTGACGTGGC). Decoy transfection data showed that transfection effciency was highest at decoy concentration of 50 nM in that cell line. Endpoints included cell number, CellTiter-Glo (Promega) and cell-cycle fractions. Differential-expression analyses used untreated target-TF expression change from 4h to 72h (single-cell RNA-seq analysis). Target Family Leverage was defined as Target CC divided by the CC of its TF family before treatment. Vehicle-treated networks and target expression were measured at 4h and 72h. N = 3 per condition.

### Core and Reactome analyses

The Core was defined independently in vehicle- and decoy-treated networks as the top 5% of TFs ranked by CC. Core replacement was calculated as the fraction of TFs in the untreated 72h Core that were absent from the corresponding treated 72h Core. Reactome pathway scores were calculated from transcriptomic data using SeuratExtend^20,24^ and compared between decoy- and matched vehicle-treated cells at 72h within each cell-cycle phase. For each of the 204 pathways, we calculated the absolute difference between treated and control scores. Overall pathway reorganization was quantified as the mean absolute score change across all pathways. The 90th percentile of absolute score changes was used to characterize the magnitude of change among the most responsive pathways. Because the direction of these score changes does not independently establish pathway activation or repression, only their magnitude was interpreted.

### Statistics

Statistical analyses were selected according to the experimental design. Analysis of variance (ANOVA) was used to compare cell-confluence responses across treatment conditions. The Friedman test was used for repeated-measures comparisons of TF CC values across matched disease states, followed by pairwise Wilcoxon signed-rank tests with Holm correction for multiple comparisons. Student’s t-test was used for comparisons of mean values between two groups. Associations between CC, network reorganization, pathway changes and cellular outcomes were evaluated using two-sided Spearman rank correlation. Sample sizes and statistical significance are reported in the text and corresponding figures and legends.

